# Pooled versus structured inhibition drives competitive stimulus interactions across distinct spatial scales of the superior colliculus

**DOI:** 10.64898/2025.12.16.694660

**Authors:** Arunima Banerjee, Ninad B. Kothari, Shreesh P. Mysore

## Abstract

Competitive stimulus interactions across the space map in the midbrain superior colliculus (SC) are essential for spatial decision-making. Here, with electrophysiological recordings in intermediate and deep layers of the mouse SC (SCid), we discovered that rules and the spatial profiles of these competitive interactions are different between local, within-RF versus global, across-RF-boundary spatial scales. When one visual stimulus was centered in the receptive field (RF) and a second one was located within the RF’s classical inhibitory surround, we found that stimulus interactions followed an averaging rule. This classical surround was spatially restricted, and the strength of inhibition underlying these ‘within-RF’ interactions decreased with distance from the RF center. By contrast, when the second stimulus was located outside the RF, stimulus interactions followed a divisive rule. Strikingly, this extra-classical inhibitory surround was spatially global, and the strength of competitive inhibition underlying these ‘across-RF-boundary’ interactions was distance-invariant. Computational modeling revealed that whereas within-RF interactions are well-explained by a divisive normalization-like mechanism driven by pooled inhibition, across-RF-boundary interactions are not, and instead, are well-explained by a winner-take-all-like mechanism driven by structured, donut-like inhibition. Combined, our results offer insights into the mechanistic logic of stimulus competition in the SCid, which critically underlies spatial decision-making in mammals.

## INTRODUCTION

Animals are exposed to a barrage of diverse inputs from multiple locations in the sensory field. Selecting the location of the most salient (or behaviorally relevant) stimulus from this array is essential for behaviors involving spatial attention, perceptual decision-making, and orienting movements. Experimental studies as well as computational models have suggested that such selection emerges from competitive interactions among the representations of stimuli in the sensory field (Desimone & Duncan 1995, Knudsen 2007, Koch & Ullman 1987). These interactions create a distributed representation of stimuli in the form of a spatial map of relative stimulus salience (or more generally, of relative priority; priority = salience + relevance) that guides the locus of selection (Fecteau & Munoz 2006, Itti & Koch 2001, Mysore & Kothari 2020).

Topographically organized maps of stimulus salience (and more generally, of priority) are known to exist in several interconnected brain regions: the frontal eye field (FEF), the lateral intraparietal area (LIP), and the superior colliculus (SC) (Goldberg et al 2006, McPeek & Keller 2004, Thompson & Bichot 2005). Among these, the superior colliculus (SC), a layered midbrain structure that is evolutionarily conserved across vertebrates, plays a central role in mediating spatial target selection (Basso & May 2017, Krauzlis et al 2013). Its intermediate and deep layers (SCid) integrate multisensory inputs into topographic representations of sensory space, and drive orienting behaviors, including saccades in primates and head movements in rodents (Allen et al 2021). Importantly, SCid activity reflects the relative priority of the stimulus, and causal perturbations in this region disrupt target selection among distractors (Lovejoy & Krauzlis 2010, McPeek & Keller 2004, Wang et al 2020).

A prominent signature of competitive selection in the SC occurs in the form of response suppression. Studies across species have shown that when multiple visual stimuli are presented simultaneously, SC responses to a given stimulus are often reduced compared to when it is presented alone (cats: (Rizzolatti et al 1973); primates: (Basso & Wurtz 1998); barn owls: (Mysore et al 2010); mice: (Wang et al 2020)). This suppression occurs for spatially distant stimuli (Mysore et al 2010, Rizzolatti et al 1974), implicating long-range competitive interactions, as well as for stimuli within a single receptive field (Alvarado et al 2007, Li & Basso 2005), suggesting local interactions. Mechanistically, lateral inhibition— the suppression of activity in one spatial channel arising from neighboring channels—is thought to play a key role in mediating these competitive interactions (Munoz & Istvan 1998, Mysore & Knudsen 2012). However, the spatial and computational principles underlying these inhibitory interactions remain poorly understood in the mammalian SC.

In this study, with extracellular neural recordings in the mouse SCid, we investigate the rules of visual stimulus competition across the SCid space map. We find that the computational rules governing paired stimulus responses, as well as the underlying spatial structure, are markedly different across two regimes: inside versus outside the classical inhibitory surround. When two stimuli fall within the RF of a neuron, the joint responses follow an averaging rule, consistent with the classical center-surround RF structure. The underlying inhibition drops off in strength with distance from the RF center. By contrast, when the competing stimulus lies outside the RF, the joint responses are suppressed with respect to the response to the RF stimulus alone, following a divisive rule. This suppression is global in scope - spanning nearly the entire azimuthal and elevational space, and its strength is space-invariant - showing no dependence on the relative distance between the stimuli. Computational modeling to account for the observed results from stimulus competition experiments reveals the necessity of two separate inhibitory circuit mechanisms operating over distinct spatial scales in order to mediate stimulus competition across the SCid space map.

## MATERIALS AND METHODS

### Animals

All mice used for electrophysiological studies were adult (8+ weeks old; 4 male and 4 female) wild-type mice (C57BL6/J strain, Jackson Laboratory). Upon arrival, all mice were housed in a colony where temperature (∼74°F) and humidity (∼50%) were controlled on a 12:12 hr light:dark cycle, and all procedures were performed after allowing mice to acclimate for at least 7 days in the new environment. All procedures followed NIH guidelines and were approved by the Johns Hopkins University Animal Care and Use Committee.

### Stereotactic surgery

Surgical procedures were followed as described previously (Kothari et al 2025). Briefly, animals were anesthetized with isoflurane (3% induction, 0.5–2% maintenance) and were secured using standard earbars in a stereotaxic frame (Kopf Instruments). A feedback-controlled heating pad (FHC) was used to maintain the body temperature at 36.8°C, and eye lubricant was applied to the eyes to prevent them from drying. Meloxicam (0.1 mg/kg) was administered as an analgesic. After the head of the animal was leveled in the stereotaxic frame, fur was gently trimmed away on top of the head, exposed skin was cleaned using betadine (antiseptic) with sterile wipes, and bupivacaine (0.75 mg/kg) was injected subcutaneously for perioperative analgesia. A scalp incision was made along the midline, followed by skin retraction, skull cleaning and preparation. Once bregma and lambda were exposed, the skull was zeroed by adjusting the earbars and bitebar of the stereotax. A craniotomy was made above the SC (centered at AP -3.5 mm; ML +/-0.8 mm from bregma). All coordinates are based on a standard mouse brain atlas (Franklin & Paxinos 2019). Craniotomies were covered with KwikSil (World Precision Instruments) for the protection of the underlying tissue. A custom-made titanium headbolt was stereotactically affixed to the skull using Metabond (Parkell), and the bregma location was kept exposed. Subsequently, mice received meloxicam (0.1 mg/kg) daily for up to 3 days.

### Neurophysiology

Neurophysiological protocols followed those that have been described previously (Kothari et al 2025). Briefly, acute single-shank silicon probes (Neuronexus 0.3 – 1.5MΩ at 1 kHz probes: A1x32-poly3-10mm-25s-177-A32, A1x32-poly3-10mm-50s-177-A32, A1x32-poly2-5mm-50s-177-A32) were used to record single and multi-units extracellularly in head-fixed, awake mice. The apparatus allowed targeting coordinates of the brain stereotactically. Neural activity that was synchronized with visual stimulus presentation times was collected via a 32-channel headstage (Intan Technologies), sampled at 30 kHz, and stored using a data acquisition system (OpenEphys).

All recordings were made in the intermediate to deep layers of the SC (SCid). Electrodes were lowered into the brain using measurements from the Bregma as well as from the pial surface. All recordings were made in the SCid by targeting depths more than 1.7 to 2 mm below the pial surface, and more than 400 μm below the surface of the SC. Recording layers/depths were verified during penetration using electrophysiological signatures and were confirmed with post hoc histology.

### Preprocessing and spike sorting

Multi-unit spike waveforms were sorted offline into putative single units using Waveclus (Quiroga et al 2004) and analyzed using custom MATLAB code. Briefly, a custom script was first used to detect and remove motion artifacts that were correlated across channels, using correlated responses evoked from potentially distinct units across channels that spiked concurrently to the stimulus. Following this, spikes were detected for individual channels at time points where the membrane potential crossed a threshold of 6-7 times the median absolute deviation. Waveforms were extracted spanning windows of 1.5 ms around the detected peak and sorted in Waveclus. To prevent detection of duplicate units across channels, only units from channels that were > 50 μm away were included for spike sorting.

### Visual stimuli

Visual stimuli were created using customized MATLAB software (Kothari et al 2025, Mahajan & Mysore 2022, Mysore et al 2010, Schryver et al 2020). Mice were head-fixed in the apparatus with the monitor 12.5 cm from the center of the eyes, in either of two positions: (#*1) Head-on/Symmetric position*: In this configuration, the tangent monitor was positioned directly ahead of the mouse so that the monitor was exposed symmetrically to both eyes; this yielded a viewing angle of ±65° azimuth and ±35° elevation (Fig. S1A); (#*2) Rotated/Asymmetric position*: In this configuration, the tangent monitor was within ±45° around a vertical axis centered between the eyes so that the monitor was more visible to one of the eyes; this provided a maximum viewing angle of -20/+110° azimuth and ±35° elevation (Fig. S1A). In all cases, positive azimuthal angles denote contralateral locations. The visual grid was corrected for distortions resulting from presenting the stimuli on a flat tangent screen. In the schematics of all experimental protocols, the tangent monitor is shown in head-on position for simplicity, even though data were collected from both positions. We confirmed that the rotation of the mouse relative to the screen did not affect the response properties of SCid neurons (Fig. S1B).

#### Expanding dots

Expanding visual stimuli consisted of a white or black dot on a gray background that increased linearly in size over the period of stimulus presentation (250 ms), starting from a size of 3° diameter. The center of an expanding stimulus remained constant throughout the duration of stimulus presentation, and expansion speed was defined as the (linear) rate of change of the angular size of the dot, *dθ*/*dt*, where *θ* is the visual half-angle subtended by the object at the eye. Different expansion speeds were achieved by changing the final size of the dot, while keeping the initial size and the duration of presentation fixed (as above). Linearly expanding dots, used extensively in previous work (Kothari et al 2025, Mysore et al 2010, Mysore et al 2011, Schryver & Mysore 2019, Schryver et al 2020), evoked stronger responses than stationary dots of the same final size (Fig. S1C), indicating that a major driver of their responses is their dynamics (speed of expansion) than simply the final size. They also evoked stronger responses for faster expansion speeds (Fig. S1C), allowing for expansion speed to serve as a metric of stimulus salience (Kothari et al 2025, Mysore et al 2010, Mysore et al 2011), and produced responses that were more resistant to trial-to-trial adaptation than stationary dots when presented repeatedly at the same spatial location (Kothari et al 2025, Mysore et al 2010). In our experiments, expansion speeds (final dot sizes) were chosen such that the responses remained within the linear range of the response curve (Fig. S1C; bottom, black curve): final diameters of the RF stimulus were 6°-10°, while those of the competitor stimulus were 12°-20°.

#### Spatial tuning curves

Spatial tuning to visual stimuli was measured by presenting expanding dots of 5° final radius, duration 250-500 ms, ISI 2.5-3.5 s, white or black, on a gray background. Tuning to visual azimuth (or elevation) was first estimated by presenting 10–15 repetitions of stimuli at randomly interleaved azimuthal (elevational) locations, with the elevation (azimuth) held constant at the manually estimated best value for the site. These measurements were iteratively performed if the azimuthal or elevation responses were different from the manually estimated centers.

#### Bar stimuli

We used bar stimuli to estimate the extent of the excitatory and inhibitory components of an SCid neuron’s classical receptive fields (Schryver & Mysore 2023). Briefly, bar stimuli were of fixed thickness (3°), but varying lengths such that the center of the bar was displayed at the receptive field center of the measured neuron in the SCid.

#### Stimulus adaptation

Responses of neurons in the SCid habituated (weakened) to the repetition of any stimulus (light, dark, moving, or stationary dots) (Kothari et al 2025, Lee et al 2020). To minimize these effects, we used long interstimulus intervals (up to 3.5 s) and used expanding dot stimuli instead of stationary dot stimuli (Kothari et al 2025, Mysore et al 2010).

#### Two-stimulus protocols

One of the main goals of this study was to study the neural responses of SCid neurons when two stimuli were present. To achieve this, visual stimuli were created following previously established protocols (Mysore et al 2010). Expanding dots were used for both stimuli. To estimate the display locations, first, the center of the receptive field (RF) was measured as described above, and a stimulus S1 was placed at the center. For two-stimulus protocols with a stimulus outside the classical receptive field, this second stimulus (S2) was typically placed >30-40° from the center. This was to ensure that S2 was sufficiently outside the region influenced by the classical surround. Post-hoc analysis ensured that S2 locations that were overlapping with the estimated RF boundary were eliminated from the analysis. When both stimuli were presented inside the RF, it was ensured that their spatial locations were non-overlapping.

### Eye tracking

To track eye positions during stimulus presentations, a camera (Arducam UC-844) was placed approximately 11 cm from the contralateral eye. Each eye was illuminated using an array of four infrared LEDs (940 nm). Video streams were collected with an average frame rate of 100 Hz and stored within a Raspberry Pi 4 module, along with TTL pulses synchronized to the onset of stimulus presentations. To extract pupil positions, a neural network was trained with four landmarks for pupil position and one for the center of the corneal reflection, using DeepLabCut (Fig. S2A, (Mathis et al 2018)). Custom MATLAB scripts were used to fit a circle to the estimated pupil coordinates and estimate the pupil centers at any instant. Pupil angular positions with respect to the eye center were calculated from the two-dimensional image using previously published methods that use the corneal reflection and a constant eyeball radius of 1.25 mm for C57BL6/J mice (Sakatani & Isa 2004). The small fraction of pupil displacements was restricted to 3.27° along the horizontal plane and 2.88° along the vertical plane (99-percentile values) (Fig. S2B,C). These results demonstrated that eye movements were small compared to the spatial extents of visual stimuli and were unlikely to modulate neural responses, particularly during two-stimulus protocols.

### Data analysis

In all experiments, the net responses of an SCid neuron to a stimulus were computed by subtracting the mean firing rate during the pre-stimulus baseline period (averaged across all interleaved trials) from the mean firing rate within a post-stimulus analysis window ([70 ms, 250 ms] w.r.to stimulus onset.

#### Receptive field (RF) center mapping

The visual RF for each site was defined as the region of the visual field where a single visual stimulus elicited a significant increase in firing rate above baseline. A location was considered to be within the RF if the baseline-subtracted response at that location was significantly greater than zero (p < 0.05, one-way ANOVA, followed by multiple comparison correction). The RF center was defined as the peak of a one-dimensional Gaussian fit to the tuning curve along azimuth or elevation.

#### Two-stimulus interactions with S2 outside the RF

To quantify competitive suppression from a second, spatially discrete stimulus (S2) *outside* the RF, we compared responses to the simultaneous presentation of S1 (centered in the RF) and S2, with responses to S1 alone (Fig. 4). A robust linear regression (*robustfit* in MATLAB) was applied to the paired response data. The intercept was considered significantly different from zero if the associated p-value was less than 0.05. The slope was considered significantly different from unity if its 95% confidence interval did not include 1.

To evaluate the effect of different locations of S2, we computed the percent change in response to S1 in the presence of S2 compared to the response to S1 alone. To identify locations where S2 had a significant modulatory effect (p < 0.05), a one-way ANOVA followed by multiple comparison correction was applied to the distribution of percent-change values. To visualize the spatial profile of suppression, we binned S2 locations into 10°-30° bins. For each bin, the mean percent suppression was calculated and plotted ± SEM. Bin centers were computed as the mid-points of the S2 locations within each bin and shown with corresponding SEM error bars.

#### Difference of Gaussians (DoG) model

To characterize classical surround suppression, we analyzed responses to bar stimuli of increasing length, which followed a classic non-monotonic profile (Fig. 5). We fit a standard difference-of-Gaussians (DoG) model, widely used in retinal and cortical physiology (Mysore et al 2010, Sceniak et al 1999), which assumes spatially co-centered excitatory and inhibitory Gaussian components that sum linearly to produce a net response *R*(*s*) described as:

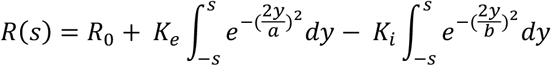

where *R*_0_ denotes the baseline response, *K*_*e*_ and *K*_*i*_ denotes the strengths, while *a* and *b* denote the space constants of the excitatory and inhibitory Gaussians, respectively. The spatial extent of the classical inhibitory surround was estimated as the bar length at which the responses reached 5% of the steady-state value. The suppression index was calculated as the difference between the peak response and the steady-state response, normalized (divided) by the peak response.

#### Time course analysis

To measure any differences in the time course of responses, we computed the instantaneous firing rates from the peristimulus time histograms (binned at 10 ms) using a Gaussian filter (σ = 12 ms). For two-stimulus experiments within the RF, we compared the instantaneous firing rates of S1 and S2 presented together with either S1 or S2 presented alone, by performing an ANOVA across each time bin. The time-to-divergence (TTD) was estimated as the first time bin where the p-value of ANOVA dropped below 0.05 and remained below 0.05 for at least 30 ms. Similarly, when the second stimulus was outside the RF, we computed the time-to-suppression (TTS) as the time point when the instantaneous firing rate in response to S1 and S2 presented together dropped significantly below the response to S1 presented alone.

#### Statistics

All statistical analyses were carried out on MATLAB. Statistical significance was accepted as p<0.05. Parametric tests (ANOVA, t-test) were used when the data were normally distributed (tested using Kolmogorov-Smirnov test), and non-parametric tests (sign test, Wilcoxon rank sum test) were performed otherwise. When testing across multiple groups, Holm-Bonferroni corrections for multiple comparisons were performed. Data is summarized as mean ± SEM unless otherwise specified.

### Modeling

#### Computational modeling of competitive stimulus interactions in SCid using spiking neurons

We used biophysically plausible neurons to simulate excitatory and inhibitory neurons for modeling. For each neuron, the membrane voltage *V*(*t*) was modeled as a first-order leaky integrator with a passive membrane time constant (Burkitt 2006, Gerstner et al 2014, Mysore & Quartz 2005).

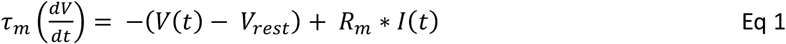

where *τ*_*m*_ is the membrane time constant, *V*_*rest*_ is the resting potential, and *R*_*m*_ is the membrane resistance. When the membrane potential reached the spike-initiation threshold, *V*_*th*_, it was clamped to *V*_*spike*_ (voltage at the peak of the spike) for one timestep, and then reset to *V*_*reset*_. A refractory period *τ*_*refractory*_ prohibited spiking for a fixed duration following each spike.

##### Neuronal model parameters selection criteria

Excitatory neuron parameters were selected to match the loom-response curves and spike rates found in SC recordings (Fig. S3A). Inhibitory neuron parameters were generally chosen to be the same as those for excitatory neurons, with the exceptions of: membrane resistance, spiking threshold and refractory periods (Gerstner et al 2014):

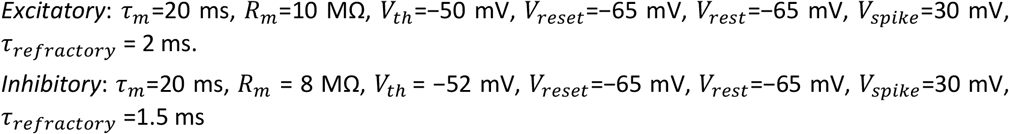

To simulate a population of neurons, we instantiated each excitatory and inhibitory neuron with randomly chosen values of the following four parameters from within the indicated ranges: *τ*_*m*_ =18-22 ms, *R*_*m*_ =8-11 MΩ, *V*_*th*_ =−52-49 mV and *τ*_*refractory*_ =1.5-3 ms.

##### Synaptic Inputs

Synaptic inputs were modeled as exponentially decaying post-synaptic currents. Every spike caused a weight-modulated (*w*_*synapse*_) step-rise in the current input to the post-synaptic neuron, which then decayed based on a time-constant *τ*_*decay*_:

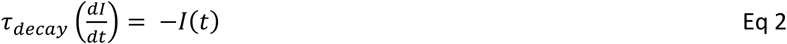

Consequently, for each spike event, a synapse would have instantaneous increments in input current as in Eq 3:

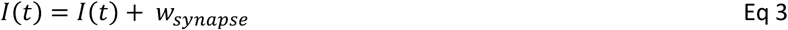

##### Synaptic weights and decay constants

The values for weights and decay constants were selected to be within the range of previously published work (Brunel 2000).

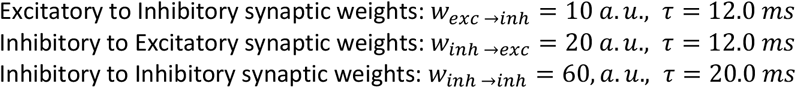

Note that the *w*_*exc* →*inh*_ weight was relevant only for the normalization circuit with pooling inhibition (Fig. 5A).

Synaptic weights were kept constant across conditions, and all excitatory post-synaptic currents contributed additively to each neuron’s net synaptic current such that the input current of a neuron ‘i’ was given by :

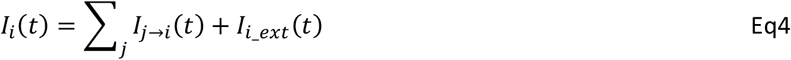

Here, *I*_*i*_*ext*_ is the external current stimulus applied to each neuron, and proportional to the strength of the stimulus (Fig. S3A)

##### Modeling the receptive field of input neurons

To model the Gaussian receptive field profile for input neurons, the external current *I*_*i*_*ext*_ was scaled by a Gaussian profile. For example, for neuron *i* with the azimuthal RF center located at position *θ*_*azim*_ and *RF width σ*_*RF*_, the effective input current with a stimulus located at *θ* was computed as:

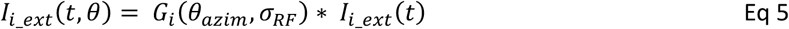

where *I*_*i*_*ext*_(*t*) is the unweighted external stimulus current, and *G*_*i*_(*θ*_*azim*_, *σ*_*RF*_) is the RF of neuron *i*:

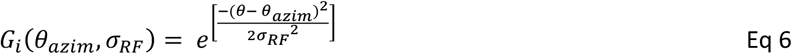

Here, *θ* is the stimulus location in azimuthal space, and *σ*_*RF*_ is the receptive-field width. This multiplicative formulation in Equation 5 allowed us to evaluate the effect of the location of both S1 and S2 on every neuron’s net input current.

##### Network circuit diagram

We modeled two circuit motifs in this study, first, the *normalization circuit with pooled inhibition* (Fig. 5A; (Mahajan & Mysore 2022, Wong & Wang 2006) and, second, the *donut-like mutual inhibition circuit* (Fig. 5C; (Mahajan & Mysore 2022, Mysore & Knudsen 2013, Wang et al 2004). Each circuit contained two input channels corresponding to the two excitatory inputs S1 and S2, each of which injected external currents (*I*_*i*_*ext*_, see Eq 4) denoted respectively as I_S1_ and I_S2_ into the input neurons. The synaptic inputs to each neuron in the models were computed as described in Equations 2 and 3.

##### Simulating competitive responses with an outside-RF competitor, using the relative-strength dependent competition protocol

For our network simulations, we adapted the competition protocol based on previously published experimental protocols in both owls and mice (Kothari et al 2025, Mysore et al 2011). Briefly, two stimuli (S1 and S2; simulated as input currents *I*_*s*1_ and *I*_*s*2_; Eqns 4 and 5) were simultaneously presented as inputs to the network. One stimulus, S1, was considered the inside RF stimulus and its strength was held constant at *I*_*i*_*extS*1_, whereas the second stimulus, S2, was presented outside the receptive field and its strength was systematically varied (*I*_*i*_*extS*2_). The resulting responses from this protocol have been termed previously as the competitor strength–response profile (CRP; (Kothari et al 2025, Mysore et al 2011)). The strength of the S2 stimulus was varied over 8-10 equal steps, ranging from below the strength of S1 to above it; S1’s strength was chosen as the middle 6^th^ or 7^th^ value in this range.

##### Analysis of simulated responses to the competition protocol

All analyses were performed using custom Matlab code. Sigmoids were fit to the responses of neuron Ex1 (Fig. 5C,E), which receives the constant strength S1 stimulus. Best sigmoidal fits to CRPs were obtained by using a nonlinear least squares estimation procedure (*nlinfit* command in MATLAB). The transition range of a CRP (i.e., precision of the neural selection boundary) was defined as the range of S2 strengths over which responses dropped from 90% to 10% of the total range of responses. The transition point of a CRP was defined as the strength of the S2 stimulus at which responses to the stimulus pair changed abruptly from high to low values, and was numerically equal to the midpoint of the transition range of the CRP. The categorization index was computed in the standard manner, per previously published work (Freedman & Assad 2006, Kothari et al 2025, Mahajan & Mysore 2022, Mysore & Knudsen 2011).

## RESULTS

### Two-stimulus interactions within the classical RF: computational rule

As a first step in investigating stimulus competition across space in the SCid, we asked what computational rules underlie short-range interactions between two stimuli occurring simultaneously within the classical RF. Studies in monkeys have shown that distinct stimuli presented inside the RF of SCid neurons interact competitively, with subsequent behavioral cues modulating this competition (Li & Basso 2005). To understand the computational rule(s) underlying such short-range competition in the mouse SCid, we examined how the responses of a neuron to a stimulus centered in its receptive field (S1) are affected by a second stimulus (S2) also located inside the receptive field. We used expanding dot stimuli as they produced robust responses compared to stationary dots of the same radius (Fig. S1C). In these experiments, S1 was presented at the center of the RF, and S2, with a faster expansion rate than S1, was presented at 2-4 locations also within the RF, always ensuring that the two stimuli did not overlap physically in space. These conditions were interleaved with trials where either S1 or S2 was presented alone at the same respective locations (Fig. 1A).

**Fig. 1.**
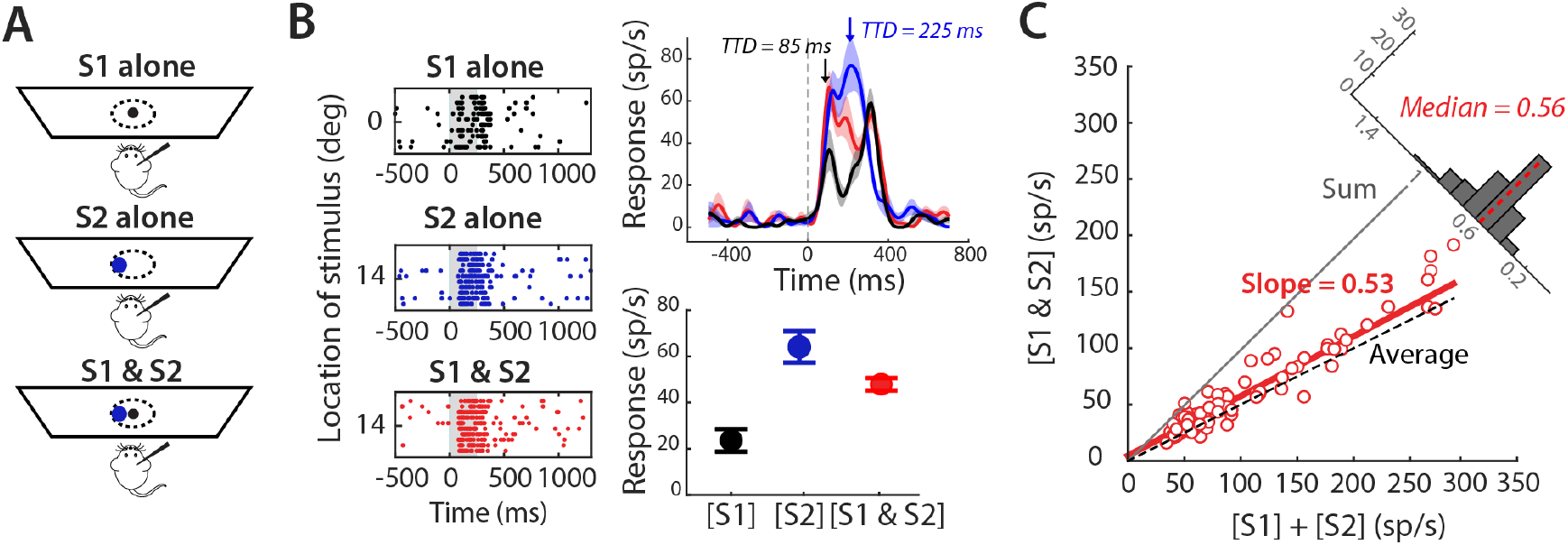
Computational rule underlying two-stimulus interactions within the RF. **A**, Schematic of the experimental setup and stimulus protocol. One visual stimulus (S1, black dot) was presented at the center of the recording neuron’s RF (dotted line), and a second stimulus (S2, blue) was presented at a non-overlapping location within the RF. **B**, Responses from an example SCid neuron. (Left) Raster plots of responses to S1 presented alone at 0° azimuth (top), S2 presented alone at 14° azimuth (middle), and both presented simultaneously (bottom). (Right, top) Average instantaneous firing rate responses to S1 alone (black), S2 alone (blue), or S1 and S2 presented together (red). Time-to-divergence (TTD, see Methods) of response to S1 and S2 presented together from S1 presented alone (black arrow), or S2 presented alone (blue arrow). (Right, bottom) Mean ± SEM of the responses corresponding to the rasters in Left. **C**, Population summary of responses to paired stimuli [S1 & S2] plotted against the sum of mean responses to individual stimuli ([S1]+[S2]) for all paired locations of S1 and S2 (n = 26 neurons, 78 paired locations). Red line indicates best-fit from a linear regression model with slope = 0.53 (not significantly different from 0.5, p = 0.107), and an intercept of 4.65 sp/s (significantly different from 0, p<0.05). (Inset) Distribution of the ratio of responses to [S1 & S2] and the sum of individual responses ([S1]+[S2]).

We observed that the responses to the simultaneous presentation of S1 and S2 were consistently lower than the linear sum of the responses to each stimulus alone, i.e., were sub-additive, and followed an averaging rule (Fig. 1B; [S1] = 23.59 sp/s, [S2] = 64.07 sp/s, [S1&S2] = 47.88 sp/s). The time course of response to [S1&S2] diverged significantly from the responses to the stronger stimulus (S2, in this example) during the late phase of the response (225 ms, Fig. 1B). Across the population, sub-additive responses were found in 77/78 of paired locations. Additionally, the averaging rule held true across the population: a linear regression model fit to the responses across all stimulus location pairs (Fig. 1C) revealed a slope of 0.53 (p = 0.107, t-test against 0.5), while the intercept was 4.65 sp/s (p = 0.035, t-test against 0).

Overall, these findings reveal that the net response to two stimuli presented within the RF of mouse SCid neurons is well-approximated by an averaging rule operating on the responses to the individual stimuli, consistent with previous reports in the primate and avian SCid (Li & Basso 2005, Mysore et al 2010).

### Two stimulus interactions across the RF boundary: the computational rule

Beyond short-range competitive interactions, SCid neurons across vertebrate species also show competitive suppression of their evoked responses by a second stimulus presented far outside the classical RF (Basso & Wurtz 1998, Mysore et al 2010, Rizzolatti et al 1974). However, the computational rules underlying such long-range interactions between distant stimuli are not known in mammals. To address this question, we examined how the responses of a neuron to a stimulus inside its RF (S1) are affected by a second stimulus (S2) outside the RF. Specifically, we presented a stimulus, S1, at different azimuthal or elevational locations spanning the RF, and the second, stronger stimulus, S2, outside the RF, at least 30-40° away from the RF center (Fig. 2A). We used expanding dot stimuli again, and fixed S2 to be stronger than S1 (expansion rates for S1: 19 ± 0.8 deg/s, S2: 43.6 ± 1.3 deg/s). Note that, by definition of the RF, S2, when presented alone, does not evoke responses different from baseline.

**Fig. 2.**
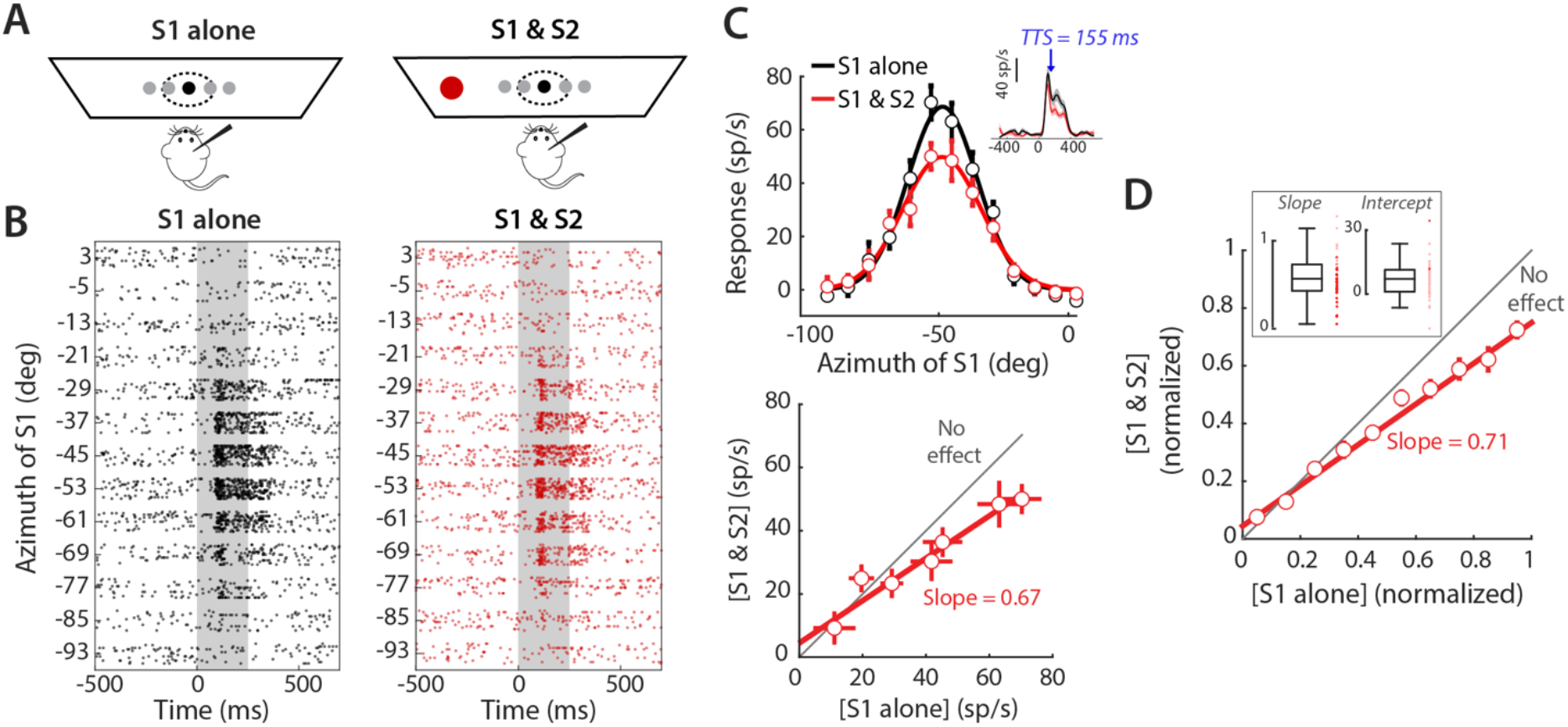
Computational rule underlying two-stimulus interactions across the RF boundary. **A**, Schematic of the experimental setup and stimulus protocol. One visual stimulus (S1, black dot) was presented at different azimuthal (or elevational) locations within the RF to measure the azimuthal (or elevational) tuning curve. S1 was presented either without (left) or with (right) a second, stronger stimulus (S2, red) at a distant, contralateral location outside the RF. **B**, Raster plots of responses to stimulus S1 presented alone at different azimuths (left) or along with stimulus S2 presented 35° away from the RF center (right). **C**, (Top) Responses of the example neuron in B plotted as a function of azimuthal location of S1, with (red) or without (black) the second stimulus, S2. Data indicate mean ± SEM. Solid lines indicate the best-fit Gaussian to the responses in each condition. (Inset) Average instantaneous firing rate responses to S1 alone (black) or S1 and S2 presented together (red). Time-to-suppression (TTS, see Methods), as indicated by the blue arrow, is 155 ms. (Bottom) Scatter plot comparing responses to S1 and S2 presented together ([S1 & S2]) against responses to S1 presented alone ([S1 alone]). Responses for only those locations at which S1 was inside the RF (i.e., at which S1 elicited above-baseline activity; see Methods) are plotted. The dashed line indicates the best-fit regression line, with a slope of 0.67, significantly different from 1 (p <0.01), and an intercept of 4.44 sp/s that is not significantly different from 0 (p =0.238). **D**, Population summary (n=40 neurons, 6 mice; 35 azimuthal tuning curves, and 5 elevational tuning curves). The responses at each neuron were normalized to its maximum response when S1 was presented alone. Data from all the neurons were binned along the x-axis (normalized responses to S1 alone; bin size = 0.1), and the average y-value (paired responses) within each group was plotted (mean ± SEM). The best fitting line to these data was determined by linear regression to have a slope (1/divisive factor) of 0.71 (significantly different from 1, p < 0.05; see Methods), and a y-intercept (additive factor) of 4% (significantly different from 0, p=0.001; see Methods).

We observed that the responses to the simultaneous presentation of S1 and S2 were lower than the responses to S1 alone (Fig. 2B,C; and therefore, also to the sum of the responses to S1 and S2). In other words, the introduction of S2 suppressed the responses to the RF stimulus (Fig. 2B,C). To quantify this sub-additive relationship, we regressed the responses in the paired condition (S1 & S2) against those to S1 alone for spatial positions of S1 within the RF. The slope of the regression line was significantly less than 1 (p < 0.05, t-test), while the intercept was not significantly different from zero (p = 0.238, Fig. 2D), indicating a divisive interaction rule. Across the population, the divisive rule was consistent, with the slope of the linear fit to the normalized data being 0.71 (1/divisive factor; p<0.05, t-test against 1; Fig. 2E), and y-intercept being 4% (p=0.001, t-test against 0; Fig. 2E); a statistically significant, but small, additive factor.

Overall, these findings revealed that the net response to two stimuli occupying locations across the RF boundary (with one inside and one outside) was well-approximated by a divisive rule operating on the responses to S1.

### Spatial extent and profile of inhibition in the classical surround

Our results, thus far, show that stimulus interactions within the RF follow an averaging rule, whereas those across the RF boundary (with one stimulus outside the RF) follow a divisive rule. However, both produced two-stimulus responses that were sub-additively related to the individual responses, suggesting the involvement of inhibitory mechanisms. Might these two types of stimulus interactions utilize a shared inhibitory mechanism? To address this question, we examined the spatial organization (extent and profile) of the inhibition underlying each.

We started by examining the spatial organization of inhibition operating within the classical RF, i.e., of the classical inhibitory surround. For this, we adopted a standard protocol used in the literature to study the classical surround: we presented bars of increasing length at the center of the RF (Fig. 3A). We found that as bar lengths increased, responses first increased to a peak value, followed by a steady decline, eventually saturating to a steady-state value for longer bars (Fig. 3B).

**Fig. 3.**
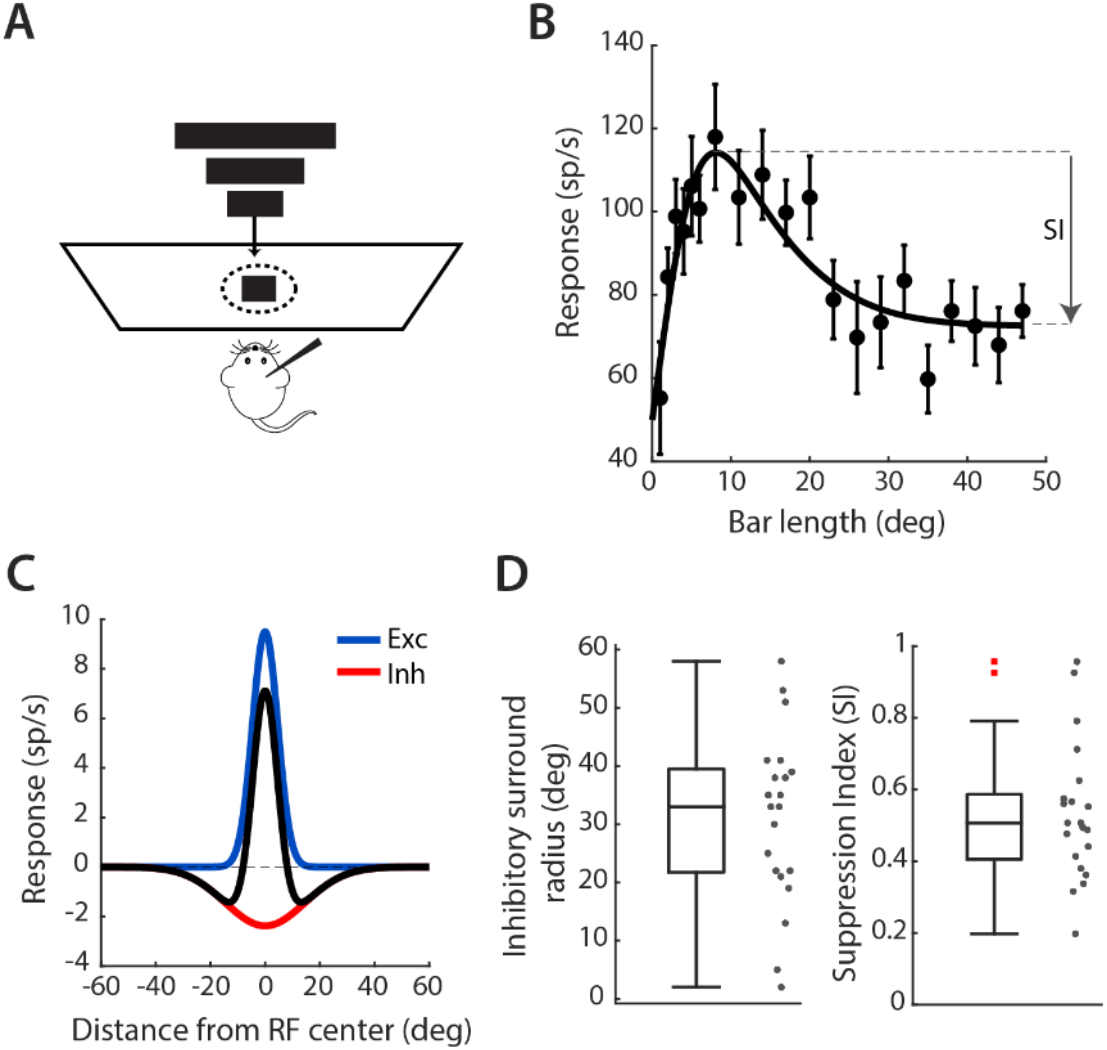
Classical inhibitory surrounds of neurons in the SCid. **A**, Schematic of experimental setup. Bars of varying lengths were presented at the center of the recording neuron’s RF. **B**, Responses of an example SCid neuron to increasing bar lengths. The solid red line indicates the best-fit line estimated using a difference of Gaussian (DoG) model shown in C. Suppression index (SI, see Methods) = 0.38. **C**, Excitatory (blue) and inhibitory (red) Gaussians obtained by fitting a DoG model to the responses in B. The net “Mexican-hat” response is shown in black. Estimated inhibitory radius for this neuron = 30° (see Methods). **D**, Boxplots and population summaries of inhibitory radii (left) and suppression indices (right) for n=22 neurons across N=3 mice. Median inhibitory surround radius = 34° and median SI value = 0.51 (p< 0.001)

The shape of such a response profile can be well-explained by a classic combination of a narrow, strong excitatory field and a spatially wider, but weaker inhibitory surround (Mysore et al 2010, Sceniak et al 1999). This would predict that for smaller bar lengths, an increasing amount of the excitatory field would get engaged as bar length increased, leading to an increase in response. For intermediate lengths, the positive effect of the spatially restricted excitatory field would be overcome by the negative effect of the wider suppressive surround, resulting in a decline in the net response. For very large bar lengths, the net contributions of the excitatory and inhibitory fields would become constant, producing a steady-state response.

To quantify this classic “center-surround” structure, we fit the data with a classic difference of Gaussians model (see Methods), consisting of an excitatory Gaussian and a wider, inhibitory Gaussian, resulting in a Mexican-hat receptive field profile (Fig. 3C). The radius of the inhibitory surround for the example site was estimated to be 30° (see Methods). Across a population of 22 neurons, the median radius was 34° (Fig. 3D). We also quantified the strength of asymptotic suppression, i.e., constant suppression at large bar lengths, using a suppression index (SI), defined as the normalized reduction of the asymptotic response from the peak response. Across the population, the median SI was 0.51 (p < 0.001, sign test against 1; Fig. 3D).

Overall, these results revealed that a spatially restrictive classical surround provides inhibition inside the RF, with its strength decreasing with distance from the RF center.

### Spatial extent and profile of inhibition in the extraclassical surround

Next, we examined the spatial organization of inhibition operating outside the classical RF. The divisive modulation of evoked responses by a stimulus outside the RF suggests a more expansive extraclassical suppressive field. To characterize the extent and spatial profile of this inhibition, we measured the effect of varying the location of a second stimulus outside the RF on the responses to an RF stimulus. Briefly, we presented an expanding dot stimulus, S1, at the center of the RF of SCid neurons, and a second, stronger, expanding dot stimulus, S2, at various locations outside the RF. We systematically varied S2’s location along the azimuthal axis, both in contralateral and ipsilateral space (relative to the recording site), while maintaining its elevation the same as that of S1 (Fig. 4A,B). In addition, we measured single-stimulus spatial tuning by presenting either S1 or S2 alone at different azimuth locations (Fig. 4C). The single-stimulus and two-stimulus conditions were randomly interleaved. For each outside-RF location of S2, we computed the percentage change in response relative to the condition when the RF stimulus, S1, was presented alone at the RF center (Fig. 4C).

**Fig. 4.**
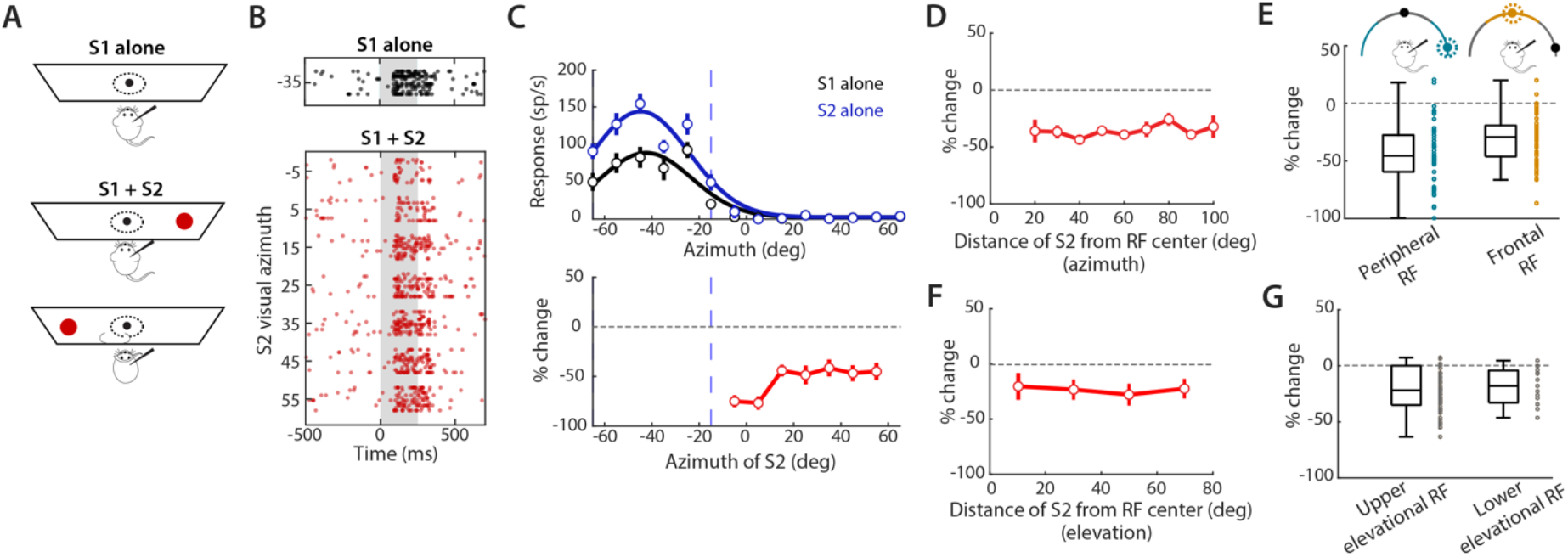
Extraclassical inhibitory surround in the SCid. **A**, Schematic of the experimental setup and stimulus protocol. In the first condition (S1 alone, black), an expanding dot stimulus (16 deg/s) was presented alone at the center of the RF. In the second, interleaved condition, S1 was accompanied by a second stimulus (52 deg/s, red) presented at different azimuthal locations outside the RF. **B**, Raster plots for an example neuron. Top: responses to S1 presented alone at the RF center (35° azimuth). Bottom: responses to paired presentation of S1 at the RF center and S2 at different azimuthal locations spanning -5° to 55° azimuth. Positive values denote contralateral location. **C**, Top: Azimuthal tuning curves for the example neuron measured independently using S1 (black) or S2 (blue). The dashed line indicates the extent of the RF (see Methods). Bottom: Percent change in responses to paired presentation of S1 and S2 compared to responses to S1 alone, plotted as a function of the location of S2 for the example neuron. Data indicate mean ± SEM. The dashed line denotes the extent of RF, estimated from the spatial tuning curve measured with S2 (top; black curve; Methods). Spatial profiles of suppression for different azimuthal locations of a competitor. **D**,**E** Population summary of extraclassical profiles of suppression for different azimuthal locations of S2 outside the RF. D, Percent change of the paired (S1 & S2) response plotted as a function of the distance of S2 from the RF center of the recording neuron. E, Percent change in response to frontal S2 stimuli for peripheral RF locations (blue), and in response to peripheral S2 stimuli for frontal RF locations (orange). All data plotted as mean ± SEM (n = 40 neurons, N = 6 mice). **F**,**G** Population summary of extraclassical profile of suppression for different elevational locations of S2 outside the RF. F, Same as D, but for different elevation locations of S2. G, Percent change in response to lower elevational S2 stimuli for upper elevation RF locations, and in response to upper elevational S2 stimuli for lower elevation RF locations. All data plotted as mean ± SEM (n = 26 neurons, N = 5 mice).

As expected, we found that the presentation of S2 produced a strong suppression in the responses to S1. Notably, we found this suppression for all locations of S2 outside the RF (Fig. 4B,C): suppression was significant at locations right outside the RF (approx. 30° from the RF center) and extended up to the tested limit of 100° (Fig. 4D).

To identify any patterns of suppression across the azimuthal space, we pooled responses from all the recorded SCid neurons in different ways. First, we examined whether suppression strength depended on the distance between S1 and S2. To this end, we aligned the RF centers of all the neurons and measured the change in response as a function of the distance of S2 from the RF center (Fig. 4D): we found that suppression strength remained constant across all distances (p = 0.279, one-way ANOVA with Holm-Bonferroni correction), with a net average suppression of 35.82%.

Next, we investigated if there were any inherent asymmetries in the suppressive interactions along the frontal→peripheral (versus peripheral→frontal) axis (Fig. 4E). To this end, we examined the suppression exerted by peripheral stimuli onto neurons with frontal RFs, and vice versa. Frontal RFs were defined as those whose centers were ≤30° in azimuth (Fig. 4D inset – orange zone), whereas peripheral RFs had centers >30° in azimuth (Fig. 4D inset – blue zone). For each group, we calculated the suppression from S2 locations in the complementary region of space. We found that neurons with frontally located RFs experienced marginally weaker suppression than those with peripherally located RFs (Fig. 4E, p<0.001, t-test).

Together, these findings reveal a widespread, effectively global, suppressive field extending across the entire azimuthal space. This extraclassical field is characterized by a lack of distance-dependence of the suppression by S2 and a consistent asymmetry in which frontal stimuli exert stronger suppression than peripheral ones.

Given the constancy and far-reaching nature of the suppression across azimuth, we next tested if these were true across elevation as well. To this end, we varied the location of S2 along the elevation axis while keeping its azimuth center at the same value as the azimuth of S1. We found no significant differences in the amount of suppression with distance from the center (Fig. 4F, p = 0.184, one-way ANOVA with Holm-Bonferroni correction). Additionally, we tested for any asymmetry in the strength of suppression from the upper versus lower elevation locations. Since elevations less than 0° are encoded by a relatively small portion of the SC (Drager and Hubel, 1976), we classified units with RF centers at an elevation of > 10° as having ‘upper’ elevational RFs, and those with RF centers <=10° as having ‘lower’ elevational RFs. In each case, we measured the amount of suppression from the complementary region of elevation space and found no significant differences between the two groups (Fig. 4G, p = 0.851, t-test).

Together, this distance-invariant, global competitive inhibition suggests the presence of an extraclassical surround mechanism distinct from that underlying the decaying-with-distance, local inhibition observed within the classical RF.

### Computational modeling of two-stimulus interactions across the SCid space map

Our experiments have revealed different computational rules and interaction profiles for the two spatial scales of stimulus interactions across the SCid space map of mice (Figs. 1 and 2). A central open question in this context is whether a common circuit mechanism underlies competitive interactions both within and outside the receptive field (RF), or whether two distinct mechanisms are needed to account for the observed results. To gain insights into potential underlying mechanisms, we turned to computational modeling (Methods). Briefly, our core network model had two ‘channels’, each corresponding to a spatial location and each consisting of excitatory and inhibitory neurons (Fig. 5). The neurons were modeled as spiking (leaky integrate-and-fire) neurons, and the values for various model parameters were grounded in published experimental findings (Methods; (Burkitt 2006, Gerstner et al 2014, Mysore & Quartz 2005)). To address questions of circuit mechanisms, we implemented different connectivity diagrams in our network model, and examined whether (and which ones) accounted for the observed experimental findings on stimulus competition.

**Fig. 5.**
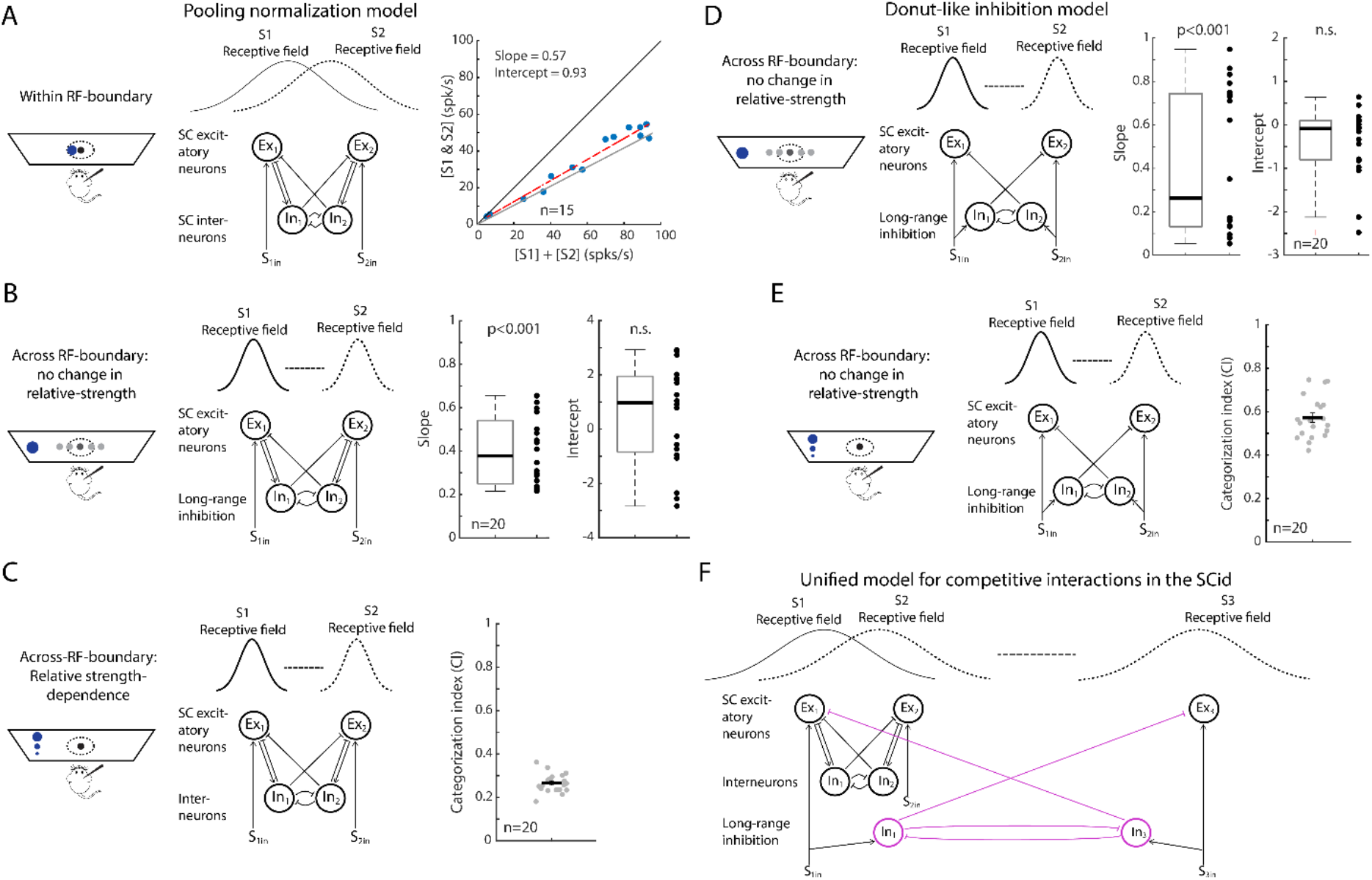
Computational modeling of two stimulus interactions across the SCid space map. **A-C**, Normalization model with pooled inhibition. (A) Within RF competitive interactions. (Left) Schematic of stimulus protocol for simulating within RF competitive interactions using the network model; these protocols are identical to those presented to the mouse (dotted outline) in Fig. 1A. (Center) Schematic of normalization model with pooled inhibition. Ex1 and Ex2 are the input excitatory neurons that receive S_1in_ and S_2in_ input stimuli. The solid and dotted line gaussian curves represent the overlapping receptive fields of Ex1 and Ex2. In1 and In2 are inhibitory neurons that send inhibition to both excitatory neurons (Methods). (Right) Population summary of responses to paired stimuli [S1 &S2] plotted against the sum of mean responses to individual stimuli ([S1]+[S2]) for all paired locations of S1 and S2 (n = 15 neurons). Red line: linear fit; gray line: represents average (p>0.9; test of slope being different from 0.5; ttest). (B) Across-RF-boundary competitive interactions. (Left) Schematic of simulation protocol for modeling across RF-boundary competitive interactions with the constant relative strength between competitors (same as Fig. 2). (Center) Schematic of normalization model with pooled inhibition. Same as in panel A, except that the receptive fields of Ex1 and Ex2 do not overlap. (Methods). (Right) Population summary of the slope and intercept (n=20 neurons; Methods; Slope; mean=0.4, SD=0.15, slope significantly different from 1, p < 0.001, t-test against 1; Intercept; mean=0.57, SD=1.87, intercept not-significantly different from 0, p=0.13, t-test against 0). (C) Across-RF-boundary interactions dependent on relative-strength: categorical responses. (Left) Schematic of simulation protocol for modeling across RF-boundary relative strength dependent competitive interactions (same as in (Kothari et al 2025)). (Center) Schematic of normalization model with pooled inhibition. Same as in panel B. (Right) Population summary of the categorization index (n=20 neurons; CI; mean=0.26, SD=0.04; Methods). **D-E**, Donut-like inhibition model (D) Outside-RF competitive interactions. (Left) Schematic of simulation protocol (same as panel B). (Center) Schematic of donut model with mutual inhibition. Ex1 and Ex2 are the input excitatory neurons and In1 and In2 are input inhibitory neurons, that receive S_1in_ and S_2in_ input stimuli. The solid and dotted line gaussian curves represent the receptive fields of Ex1, Ex2, In1 and In2. The critical difference between the donut model and normalization models is that there is no self-inhibition in the donut model (i.e. In1 and In2 do not inhibit Ex1 and Ex2, respectively; Methods). (Right) Population summary of the slope and intercept (n=20 neurons; Methods; Slope; mean=0.42, SD=0.33,, slope significantly different from 1, p < 0.001, t-test against 1; Intercept; mean= -0.38, SD=0.79, intercept not-significantly different from 0, p=0.078, t-test against 0). (E) Across-RF-boundary interactions dependent on relative-strength: categorical responses. (Left) Schematic of simulation protocol (same as panel C). (Center) Schematic of donut model with mutual inhibition. Same as in panel D. (Right) Population summary of the categorization index (n=20 neurons; CI; mean=0.57, SD=0.09; Methods). **F**, Schematic of the unified model of competitive interactions across the SCid space map involving two distinct mechanisms operating on the short, within-RF spatial scale, versus the longer range, global, across-RF-boundary spatial scale. For simplicity, this model has three input channels (S_1in_, S_2in_ and S_3in_). The normalization model (black lines) operates on the local (within-RF) spatial scale: between Ex1 and Ex2 excitatory neurons, with In1 and In2 representing the pooled inhibition. The donut-like inhibition model (pink lines) operates on the global (across-RF-boundary) scale: between Ex1 and Ex3, with In1 and In2 implementing self-sparing, mutual inhibition.

As a first step, we first considered our experimental results showing that competitive interactions within the RF follow the averaging rule. These results are consistent with prior reports of response averaging across vertebrate species, including the primate SCid (Li & Basso 2005), bird OTid (Mysore et al 2010), and primate visual cortex (Reynolds et al 1999). Because such averaging in the visual cortex has been previously modeled using divisive normalization circuits (Reynolds et al 1999), we tested whether our SCid data in mice (Fig. 1) could also be explained by a normalization mechanism. We implemented a well-established circuit diagram for divisive normalization model with pooled inhibition in our network (Fig. 5A; Methods; (Mahajan & Mysore 2022, Wong & Wang 2006)).

To simulate within-RF stimulus interactions in our model (Fig. 5A left; Fig. 1A), we set the spatial RFs of the neurons in each channel to overlap significantly with those of the neurons in the other channel (Fig. 5A, center). Channel 1 was designated the ‘recording’ channel, and S1 was ‘presented’ at the center of the RF of its excitatory neuron (Ex1). S2 was ‘presented’ to channel 2, which encoded an adjacent location to S1. Notably, by virtue of our design choice of overlapping RFs between the two channels, S2 was also located within the RF of Ex1 (the neuron being recorded). When S1 and S2 were ‘presented’, respectively, to each of the two channels, thus mimicking the within-RF experiment (Fig. 1A), we found that the responses of the ‘recorded’ model neuron, Ex1, closely reproduced within-RF experimental results from Fig. 1 (Fig. 5A). Specifically, Ex1’s responses to simultaneous S1 & S2 presentation were consistently lower than the sum of the individual responses, and followed an averaging rule (Fig. S3B, example neuron; Fig. 5A, right; population summary).

We next tested whether the same normalization model could also account for two-stimulus interactions **across the boundary of the RF** (Fig. 5B, left). To do so, we set the spatial RFs of the neurons in the two channels to be far away from each other (i.e., no spatial overlap; Fig. 5B, center). We found, again, that the responses of Ex1 closely reproduced outside-RF experimental results from Fig. 2 (Fig. 5B). Specifically, introducing S2 outside the RF caused suppression of Ex1’s responses to the RF stimulus, and followed a divisive rule (Fig. S3 C: example neuron; Fig. 5B: population summary of the slope and intercept). These results (Figs. 5AB) indicate that a classic, pooled normalization circuit **could**, in principle, serve as a common mechanism for competitive interactions both within and across the boundary of the RF of SCid neurons, in line with previous suggestions (Basso & May 2017).

However, if normalization is the common mechanism governing both spatial scales of competition, the same circuit should also explain another recent finding about across-RF-boundary competitive interactions in the mouse SCid (Kothari et al 2025). In that experiment, responses of SCid neurons were measured to a fixed-strength stimulus inside the RF (S1) while simultaneously presenting a second, competing stimulus (S2) of varying strength outside the RF. Using this competition protocol that tested the encoding of relative stimulus strength, SCid neurons were shown to signal categorically the strongest stimulus across the SCid space map. Here, we tested if these results as well could be accounted for by the normalization model. We did so by presenting to our model, S1 of fixed strength, and a distant S2 of varying strength (Fig. 5C, left).

We found that the normalization circuit model **failed to produce categorical signaling**. Responses of the SCid RF neuron (Ex1; Fig. 5C center) decreased **gradually** with increasing S2 strength, rather than exhibiting an abrupt, categorical drop (Fig. S3 E: example neuron). To quantify the categoricalness of the response profile, we computed the categorization index (CI; (Freedman & Assad 2006, Kothari et al 2025, Mahajan & Mysore 2022); Methods). The categorization index (CI) was low (Fig. 5C right: CI population summary; Fig. S3 E: example neuron), demonstrating that pooled normalization cannot reliably categorically discriminate the strongest stimulus from *other* weaker competitors. These results agree with prior modeling of experimental data from avians, which also showed that normalization with pooled inhibition is **ineffective** for producing categorical selection (Mahajan & Mysore 2022). Thus, the normalization model **cannot** account for a key finding on across-RF-boundary competition, indicating that it cannot serve as a common mechanism for within as well as across RF boundary stimulus interactions.

What, then, might be a viable mechanism for long-range, across-RF-boundary stimulus interactions in the mouse SCid? Previous work in avians provides a potential avenue of investigation. A specialized, donut-like inhibitory motif between the avian SC (called optic tectum, OT) and a satellite inhibitory nucleus called the Imc (nucleus isthmus pars magnocellularis; (Mysore & Knudsen 2013) has been identified as being critical for categorical stimulus selection by OTid neurons in avians. The mouse analog of the avian Imc, called PLTi (parabigeminal lateral tegmental inhibitory complex), was identified recently, shown to be connected bidirectionally with the SC, and was demonstrated to be involved in target selection for spatial attention in freely behaving mice (Kothari et al 2025). We, therefore, decided to test whether this circuit motif can account for both key aspects of across-RF-boundary stimulus interactions observed above in the mouse SCid, namely: the divisive inhibitory rule due to a fixed strength S2 (Fig. 2; Fig. 5D left), and the categorical response profile due to a varying strength S2 (Kothari et al 2025).

We first tested the donut-like inhibitory motif’s ability to account for the observed divisive suppression (Fig. 5D center). We found that, similar to the normalization circuit, the donut-like circuit successfully reproduced divisive suppression of S1 responses by an outside-RF S2, matching the experimental results in Fig. 2 (Fig. S3D: example neuron; Fig. 5D right: population summary of the slope and intercept). We then tested the donut-like motif’s ability to account for the relative-strength dependent responses (Fig. 5E left and center). We found that, unlike the normalization model, it successfully also generated **categorical, switch-like responses** when the strength of an outside-RF S2 was varied systematically (Fig. S3E: example neuron ). The categorization index of these response profiles was high (Fig. 5E right: CI population summary; Fig. S3E: example neuron), revealing the ability of this circuit to robustly discriminate the strongest stimulus from weaker competitors. These findings indicate that a donut-like mutual inhibition circuit can account for both the key signatures of stimulus competition in the SCid involving a second stimulus that lies outside the RF.

As a final step, we wondered whether this structured, donut-like inhibitory circuit might also be able to account for within-RF interactions, thereby serving as a potential common model for all two-stimulus interactions across the SCid space map. To address this question, we considered the anatomical connectivity of the donut-like inhibitory motif: it is defined by the self-sparing pattern of spatial inhibition that it implements. That is, the inhibitory neuron of each spatial channel delivers stimulus-evoked inhibition to all other spatial channels except itself. By contrast, the construction of the classical inhibitory surround, which governs the within-RF interactions, is driven by inhibition which peaks at the center of the spatial channel and decays with distance from it, as revealed by the bar-length experiments (Fig. 3). In other words, the inhibitory mechanism for within-RF interactions necessarily involves each spatial channel inhibiting itself. Consequently, the donut-like inhibitory motif is fundamentally incompatible with, and not a viable mechanism for, within-RF interactions.

Combined, our results indicate that two distinct inhibitory circuit mechanisms govern stimulus interactions across the SCid space map: 1) a pooled normalization-like mechanism for within-RF stimulus competition (Fig. 5F), but 2) a distinct, donut-like inhibitory mechanism for long-range, across-RF-boundary competition (Fig. 5F).

## Discussion

In this study, we investigated the principles underlying competitive stimulus interactions across two spatial scales in the SCid, namely those governed by a classical inhibitory surround and those governed by an extraclassical inhibitory surround.

### Computations underlying stimulus interactions within the classical and the extraclassical surrounds

Computationally, stimulus interactions within the RF as well as across the RF boundary were sub-additive, signaling the involvement of inhibitory influences (Basso & May 2017). However, the nature of these sub-additive interactions turned out to be different. Within the classical surround, stimulus interactions were best described by an averaging rule, where responses to two stimuli approximated the mean of their individual responses. Sub-additive interactions of this kind have been reported in the primate SC (Li & Basso 2005), the barn owl optic tectum (Mysore et al 2010), and cortical areas such as V4 (Reynolds et al 1999). Such averaging is consistent with normalization models in which responses are balanced by the pooled inhibitory activity driven by all the inputs (Carandini et al 2005, Carandini & Heeger 2011, Reynolds & Heeger 2009).

By contrast, interactions with competitors outside the RF followed a divisive suppression rule of the responses to the RF stimulus. These interactions across the RF boundary that do not involve changes in relative strength, seemed, on the surface, to be well accounted for by normalization mechanisms: both by our modeling results (Fig. 5B), and suggested by past experimental work (Basso & Wurtz 1998, Vokoun et al 2014). However, the inclusion of two-stimulus interactions involving relative strength-dependent response profiles revealed a different picture. SCid neurons across vertebrate species have been shown recently to exhibit categorical response profiles, switching abruptly from one stimulus representation to another as relative salience (or priority) of the competing stimuli changes (Kothari et al 2025, Mysore et al 2011, Mysore & Knudsen 2014). Such categorical response profiles aid winner-take-all selection among distant competing stimuli across the SCid space map, especially when the relative salience or priority of competing stimuli are close to one another (Kothari et al 2025, Mahajan & Mysore 2022, McPeek & Keller 2004, Mysore et al 2011, Mysore & Knudsen 2014), and are critical for the function of SCid (Kothari et al 2025, Lovejoy & Krauzlis 2010, McPeek & Keller 2004, Mysore & Kothari 2020). These categorical dynamics, revealed only with stimulus protocols exploring relative strength dependence, however, could not be accounted for by standard pooling normalization models (Fig. 5C). Instead, they required a self-sparing inhibition motif (Fig. 5D-F), where inhibitory neurons driven by one stimulus suppress the representations of the competing stimuli but not their own; a finding that is congruent with the recent discovery of self-sparing inhibition as the circuit mechanism underlying categorical responses in the OTid (Mahajan & Mysore 2022). Such a motif, incidentally, also successfully implements some of the classic functions ascribed typically to pooled normalization mechanisms, such as maintaining responses within the dynamic range through response suppression (Mahajan & Mysore 2022). Taken together, our results reveal the necessity of distinct mechanisms and separate neural circuits for mediating local versus global competitive interactions in the SCid.

### Classical versus extraclassical surrounds in visual processing and stimulus selection

We show that the classical and extraclassical surrounds in SCid exhibit fundamentally different properties. Classical surrounds are spatially localized, with the strength of inhibition decreasing as a function of distance from the center. Typically, such surrounds are thought to shape responses to single stimuli, such as controlling receptive field sizes and determining tuning properties to features (Ito et al 2017). However, when multiple stimuli are presented in the RF, the resulting responses could be either integrative or competitive: cross-modal interactions within the RF often produce the former (with additive or superadditive effects), while unimodal interactions often produce the latter (yielding sub-additive effects). Considering that neurons in the SCid primarily represent the salience or relevance of stimuli relative to that of others in space, i.e., that they play a major role in relative priority computations, it is reasonable to assume that sub-additive interactions between spatially resolvable stimuli of the same modality in the SC are competitive rather than integrative, even when they are presented within the same RF (Alvarado et al 2007). Indeed, experiments in the primate SC have shown that cueing one of two simultaneously presented stimuli as the target for a subsequent saccade can modulate the neural responses from a simple average to a winner-take-all-like representation of the target stimulus (Li & Basso 2005), just as in cortical areas (Reynolds et al 1999). Thus, classical inhibitory surrounds may provide the substrate for both integration (across modalities) and short-range competition (within modalities), allowing local networks to flexibly represent either multiple nearby stimuli or only the most behaviorally relevant one.

By contrast, extraclassical surrounds are spatially expansive and distance-invariant, enabling global suppression across the visual field. In addition, these competitive interactions also operate between locations encoded by opposite hemispheres. The presence of such global, intra- and cross-hemispheric inhibitory surrounds has been reported previously in the cat SC (Rizzolatti et al 1973, Rizzolatti et al 1974) and in the owl optic tectum (OT; (Mysore et al 2010)). Here, we also found that frontal stimuli exert marginally but significantly stronger suppression on peripheral representations, than vice-versa. Such an asymmetry might be ethologically relevant to rodents for prioritizing threatening stimuli occurring in the frontal space. Overall, the observed pattern of global inhibition is consistent with the requirements for the construction of a map of relative stimulus strengths (or priorities) in which stimulus strengths (priorities) across all pairs of spatial locations need to be evaluated to determine the location of the highest priority stimulus (Knudsen 2018, Mahajan & Mysore 2018).

### Circuit mechanisms for local versus global inhibition

How might these surrounds be constructed in the SCid? The SC contains a large complement of GABAergic neurons (Mize & Luo 1992). Spatially localized inhibition consistent with that required for the construction of a classical surround has been reported in the intermediate layers of the SC in rodent slices (Lee & Hall 2006, Phongphanphanee et al 2011). Functionally, local GABAergic inhibition in the SCid has been demonstrated to play a role in visually guided behavior. For instance, activation of local GABAergic inhibition impairs behavioral performance in an orientation change detection task (Hu et al 2019, Wang et al 2020). Thus, the source of inhibition for shaping local (within RF) stimulus interactions likely originates within the SCid. However, the extent to which such short-range interactions are produced by sources of inhibition within the SC versus are inherited from earlier processing stations remains to be worked out.

The SCid, however, lacks extensive, long-range inhibitory projections required for the construction of a global surround (Lee & Hall 2006). Additionally, intercollicular projections in mice are reported to be patchy and restricted to specific spatial zones rather than globally uniform: studies have reported rostro-caudal asymmetry (Bayguinov et al 2015) or exclusive rostral-rostral or caudal-caudal connectivity patterns (Doykos et al 2020). Together, these findings suggest that sources that mediate extraclassical inhibition originate largely outside the SCid.

What sources extrinsic to the SC potentially fit the bill? Although long-range competitive interactions have been previously reported in the retina (Baccus 2007, Lee 1996), these cannot explain inhibition resulting from the opposite hemisphere or from multisensory stimuli (as shown also in birds; (Mysore et al 2010)). Similarly, inputs from the substantia nigra are also unlikely to serve as the source for global competitive inhibition in the SCid as they function by a focal, stimuli-driven disinhibition mechanism that is suited for gating rather than broad suppression (Hikosaka & Wurtz 1985). In our recent work, we identified another candidate inhibitory nucleus called the PLTi, situated in the brainstem tegmentum, that provides widespread inhibition to the SC (Kothari et al 2025). Bilateral silencing of this nucleus disrupts target selection in a spatial attention task, without any impairments to motor movements. In previous works, the avian homolog of the PLTi, called the Imc, has been demonstrated not only as a source of competitive suppression in the owl optic tectum (Mysore & Knudsen 2013) but also as an active substrate for the construction of extraclassical surrounds (Schryver & Mysore 2023). Furthermore, our results, here, from bar-length tuning experiments show that broad, symmetrical activation of the extraclassical surround of SCid neurons does not produce unlimited suppression, with suppression plateauing out at a non-zero value. These observations suggest the presence of reciprocal inhibition among the inhibitory neurons mediating the extraclassical surround (Mysore et al 2010, Mysore & Knudsen 2013); indeed, such reciprocal inhibition has been found within the avian Imc (Goddard et al 2014, Mysore & Knudsen 2012). Together, these converging pieces of evidence suggest that local SCid circuits mediate classical surround suppression, whereas extrinsic, long-range sources - likely the PLTi - implement extraclassical suppression and generate the global inhibitory field needed for distance-invariant competition. Investigating the molecular identity and microcircuit connectivity of the local inhibitory neurons in the SC that may help construct the classical inhibitory surround, as well as the role of PLTi in constructing the extraclassical surround of SCid neurons are two exciting directions of future work.

